# From Prompt to Pipeline: A Comparative Evaluation of Large Language Model Coding Agents for Reproducible Bioinformatics Pipeline Construction

**DOI:** 10.64898/2026.09.11.751004

**Authors:** Paul R. Munn, Jennifer K. Grenier

## Abstract

Agentic coding systems are increasingly presented as a way to reduce the engineering burden of scientific software development. Bioinformatics is a strong test case for this claim because useful pipelines must combine domain-specific analysis choices, command-line software, sample metadata, workflow orchestration, container or HPC execution, and interpretable quality-control reporting. We evaluated three agentic systems - Biomni, Claude Code, and Codex - on the same task: constructing a Nextflow DSL2 pipeline for paired-end CUT&Tag data that included read QC, trimming, alignment, filtering, duplicate removal, signal track generation, per-sample and group-level peak calling, control-aware group merging, annotation, FRiP calculation, deepTools visualizations, and final MultiQC reporting. Each system received the same detailed CRAFT-style prompt and was assessed against a hand-coded reference pipeline developed by the authors.

All three systems produced pipeline implementations that appeared plausible at the level of documentation and file structure, but none fully satisfied the requested analysis. The most consequential failure was shared: the agent-generated pipelines performed some form of group-level merging but did not produce the requested merged-group reporting outputs. Sample-level MultiQC reports also disagreed with the reference report. Codex was closest to the reference for primary mapped-read counts, although its total-read accounting and report structure still differed. Claude Code produced the broadest final report, but its mapping summary mixed stages and therefore could not be treated as numerically correct. Biomni produced the strongest subjective documentation, but its read-count agreement with the reference report was poor and several failures required substantial Nextflow expertise to diagnose. These results suggest that current coding agents can accelerate scaffolding, documentation, and routine implementation, but they do not eliminate the need for expert review in bioinformatics workflow construction. For complex sequencing workflows, prompts must specify not only the biological intent, but also the exact stage semantics, acceptance tests, metadata contracts, expected report sections, resource propagation rules, and failure criteria needed to distinguish a plausible pipeline from a correct one.

**Author summary:** Modern AI coding agents can write large amounts of software from natural language instructions. This study asks whether that ability is enough to build a real bioinformatics pipeline for CUT&Tag sequencing data. The answer from this evaluation is mixed. The agents produced impressive-looking pipelines and documentation, but the outputs did not fully match a hand-coded reference workflow. Most importantly, the pipelines did not generate the requested merged-group reports, which were central to the biological use case. Some errors were simple programming issues, but others required a working knowledge of Nextflow, sequencing QC, and chromatin profiling workflows. The practical conclusion is not that these tools are useless. Rather, they are best viewed as accelerators for experienced users. A biology student with little programming or workflow-management background would still have difficulty detecting and fixing the most important errors.

## Introduction

Reproducible bioinformatics pipelines occupy an awkward middle ground between research software engineering and experimental biology. A useful workflow must be scientifically appropriate, computationally robust, portable across local and HPC environments, and understandable to researchers who may not be professional software developers. Workflow systems such as Nextflow have improved reproducibility by making data dependencies, parallel execution, and software environments more explicit (Di Tommaso et al., 2017). Reporting tools such as MultiQC have similarly improved large-scale quality assessment by consolidating logs and metrics across many command-line tools (Ewels et al., 2016). However, building and maintaining a specialized workflow remains a substantial burden, especially when the analysis requires complex sample metadata, biological grouping, matched controls, and custom reporting.

CUT&Tag provides a useful stress test for agentic software development. The assay profiles chromatin-associated proteins and histone modifications using antibody-targeted tagmentation, producing libraries that are often analyzed with tools originally developed for ChIP-seq (Yu et al., 2015), CUT&RUN (Kaya-Okur et al., 2019), or ATAC-seq (Grandi et al., 2022). A routine pipeline may include adapter trimming, paired-end alignment, mitochondrial and blacklist filtering, duplicate handling, peak calling, signal track generation, peak annotation, and enrichment visualization. In practice, laboratories often require additional logic that is not captured by generic examples: samples may be split by species, antibody, condition, replicate, and control relationship; per-sample peaks may need to be distinguished from merged group-level peaks; and final reports may need to summarize both individual libraries and biologically meaningful merged groups. These requirements are conceptually straightforward but fragile in implementation.

Large language model coding agents now promise to automate many of the tasks involved in producing such software. Codex was introduced by OpenAI as a cloud-based software engineering agent capable of reading a codebase, editing files, running commands, and proposing changes for review (OpenAI, 2025). Claude Code is described by Anthropic as an agentic coding tool that works from the terminal, IDE, or GitHub and can understand and edit codebases through natural language commands (Anthropic, 2026). Biomni is a biomedical agent designed to combine language-model reasoning, biomedical tool selection, retrieval, and executable code construction across biomedical tasks (Huang et al., 2025). These systems differ in interface and emphasis, but all raise a similar practical question: can a researcher describe a complex biological workflow in detailed natural language and obtain a working, trustworthy pipeline?

This paper evaluates that question using a deliberately realistic task. Three agentic systems - Biomni, Claude Code, and Codex - were asked to build a production-quality Nextflow DSL2 pipeline for paired-end CUT&Tag FASTQ processing (**Figure 1**). The prompt specified required inputs, sample association metadata, read pairing, trimming, alignment with Bowtie2, filtering, duplicate removal with Picard, optional UMI handling, BigWig and BedGraph track generation, MACS3 peak calling, group-level peak merging, matched control handling, annotation, FRiP calculation, deepTools visualizations, and multiple MultiQC reports. The generated pipelines were then assessed in terms of documentation, setup burden, bugs encountered, technical expertise required for debugging, and agreement with the author’s hand-coded reference pipeline.

**Figure 1.**
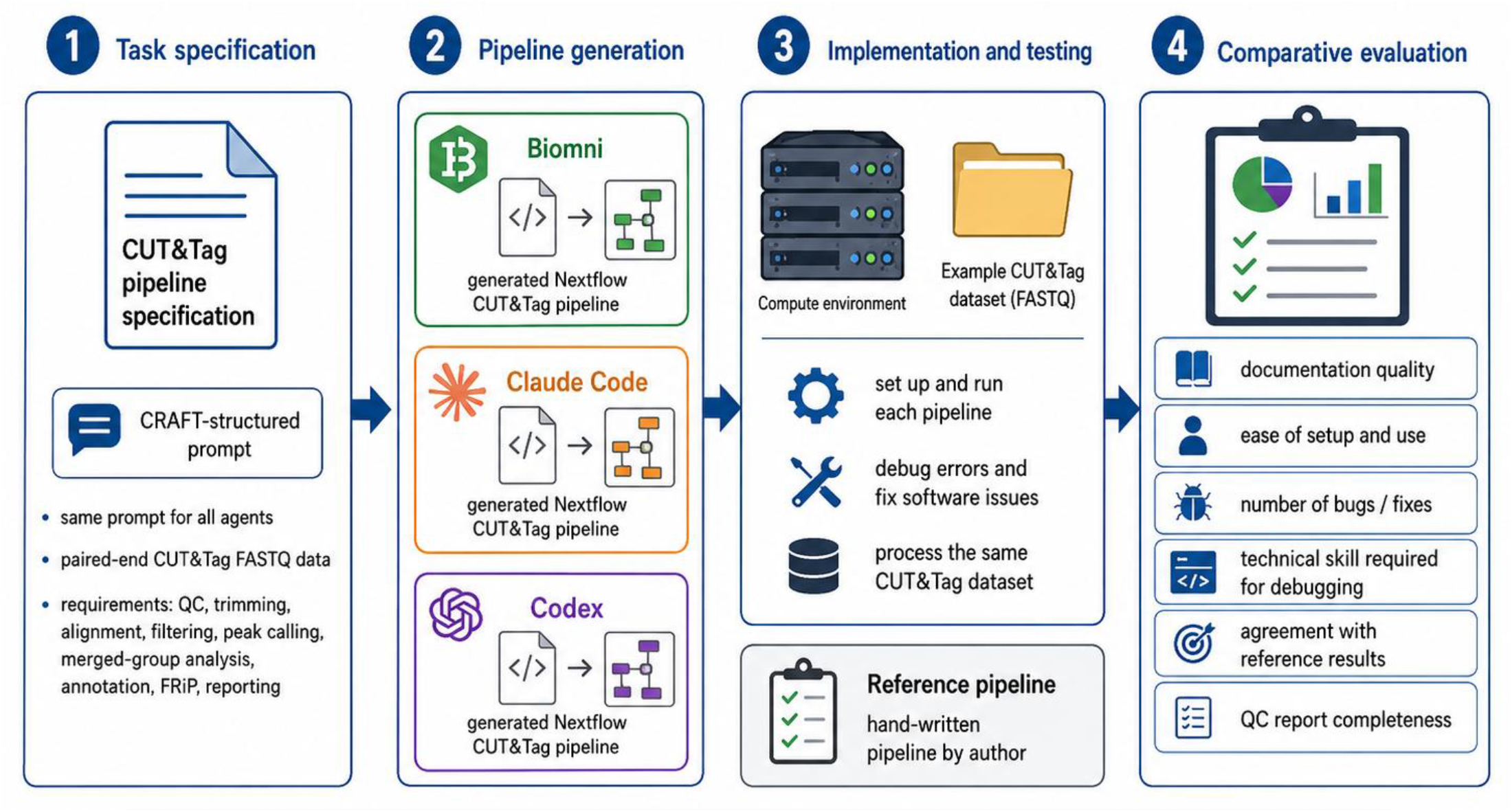
Study design for evaluating agentic coding systems in bioinformatics pipeline construction. A single detailed CUT&Tag pipeline specification was provided independently to three agentic coding systems: Biomni, Claude Code, and Codex. Each system was tasked with generating a reproducible Nextflow pipeline for paired-end CUT&Tag data processing, including FASTQ discovery, quality control, trimming, alignment, filtering, duplicate removal, peak calling, group-level merging, annotation, FRiP calculation, signal-track generation, and MultiQC reporting. The resulting AI-generated pipelines were then run on the same input FASTQ files and compared against a hand-written reference pipeline. Evaluation focused on documentation quality, ease of setup, number and severity of bugs, technical expertise required for debugging, agreement of read-count and QC metrics, and completeness of final biological reporting, especially for merged sample groups.

The central finding is that agent-generated pipelines were useful but not reliable. All three agents produced credible implementations, and in some cases substantial documentation. Yet none fully reproduced the requested analysis outputs. The most important failures were not superficial style problems. They concerned stage definitions, Nextflow channel semantics, control-aware grouping, report generation, and interpretation of QC metrics. These are exactly the parts of a scientific workflow where a plausible implementation can produce misleading results if not carefully audited.

## Results

### Benchmark task and evaluation design

The benchmark consisted of a single complex pipeline-construction task. Each agent received the same specification, written using a CRAFT-style structure: context, role, action, format, and target audience (see Methods). The specification asked for a complete Nextflow DSL2 implementation and documentation package, not just a conceptual outline (**Figure 2, Supplemental File 1**). It required the workflow to scan nested input directories for paired-end FASTQ files, validate R1/R2 pairing, match sample IDs to an association CSV, restrict processing to samples matching the requested genome and fail early when associations were missing or inconsistent. Downstream steps included raw and trimmed FastQC, cutadapt trimming, Bowtie2 alignment, samtools statistics, mitochondrial and blacklist filtering, Picard duplicate removal, optional UMI-aware deduplication, read-retention reporting, deepTools signal tracks, per-sample and merged-group MACS3 peak calling, peak annotation, FRiP calculation, TSS and peak-centered visualizations, and final MultiQC reporting.

**Figure 2.**
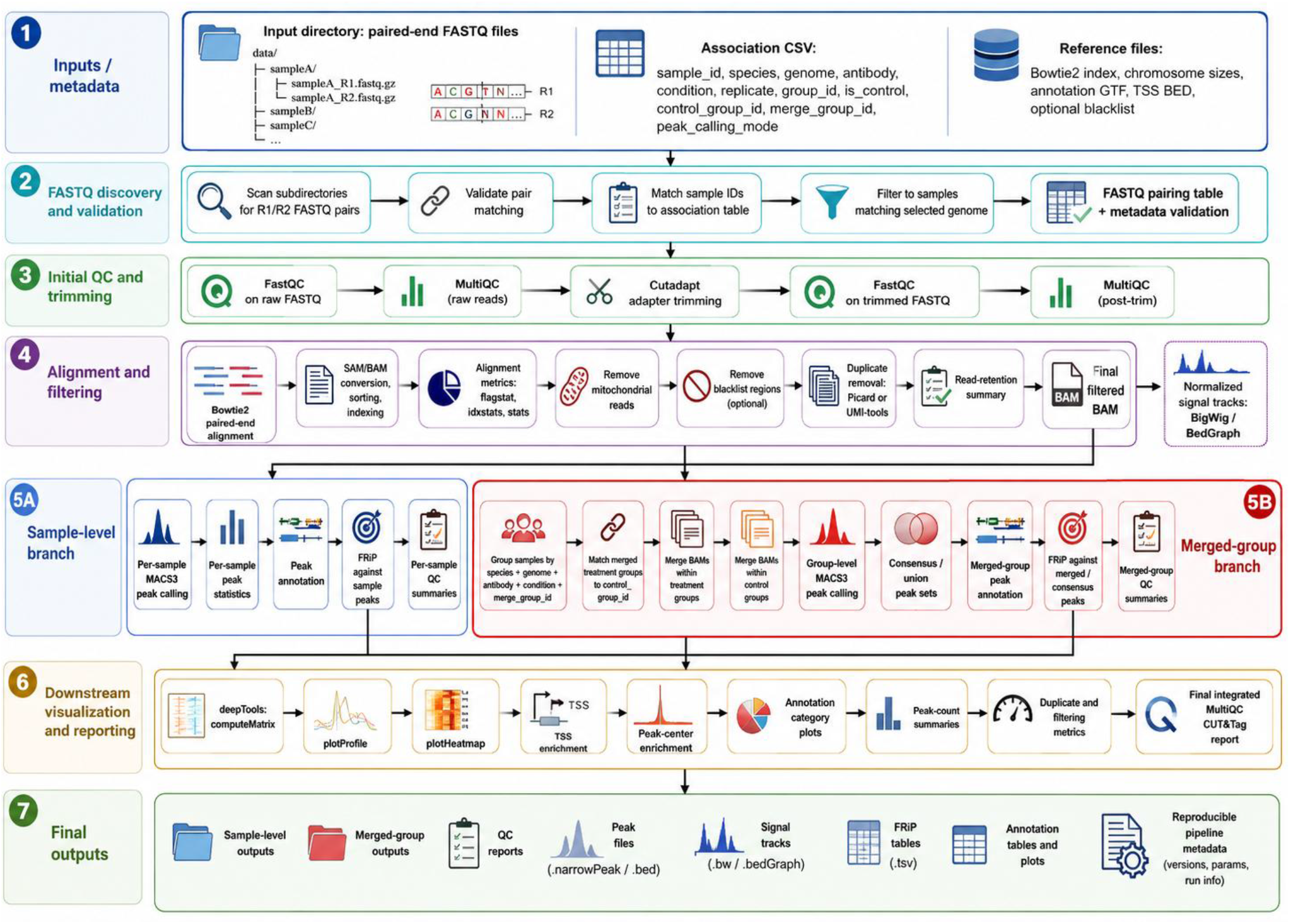
Requested CUT&Tag pipeline architecture. The requested workflow was designed as a full paired-end CUT&Tag processing pipeline beginning with recursive FASTQ discovery, R1/R2 pair validation, and matching of sample identifiers to an association table containing species, genome, antibody, replicate, control, and merge-group metadata. After initial raw-read QC, reads are adapter-trimmed, re-assessed by FastQC/MultiQC, aligned to the selected reference genome with Bowtie2, sorted and indexed, and filtered to remove mitochondrial reads, blacklist-overlapping reads, and duplicate fragments. The filtered sample-level BAM files are then used to generate normalized BigWig and BedGraph signal tracks. The architecture includes two major analysis branches: a **sample-level branch**, in which MACS3 peak calling, peak annotation, FRiP calculation, and QC summaries are generated for each individual sample; and a **merged-group branch**, in which samples sharing a defined `merge_group_id` are merged, matched to the appropriate merged control group, subjected to group-level peak calling, and used to generate consensus peak sets. Downstream reporting integrates deepTools enrichment profiles, heatmaps, TSS enrichment, peak-center enrichment, annotation-category summaries, peak counts, duplicate and filtering metrics, FRiP tables, and final MultiQC reports, producing both sample-level and merged-group outputs with reproducibility metadata.

The reference workflow used a conventional chromatin-profiling toolchain. Adapter trimming was performed with cutadapt, alignment with Bowtie2, alignment manipulation and reporting with samtools, duplicate handling with Picard, peak calling with MACS3, read-in-peak quantification with featureCounts, and peak annotation with HOMER. These choices are broadly consistent with published or widely used tools for short-read chromatin-profiling workflows: cutadapt for adapter removal (Martin, 2011), Bowtie2 for paired-end alignment (Langmead and Salzberg, 2012), SAMtools for alignment processing (Danecek et al., 2021), MACS for enrichment-based peak calling (Zhang et al., 2008), featureCounts for read summarization (Liao et al., 2014), HOMER for motif and genomic annotation workflows (Heinz et al., 2010), and deepTools for signal enrichment plots and heatmaps (Ramirez et al., 2016).

### Documentation quality differed from executable correctness

All three agents produced documentation that was at least superficially useful, but documentation quality did not predict correctness. Biomni produced the strongest documentation subjectively. Its output was broad, domain-aware, and close in spirit to the requested pipeline blueprint. Claude Code produced the second strongest documentation and adopted an approach similar to Biomni, which was notable because the two systems are architecturally and operationally different. Codex documentation was useful but less comprehensive than the other two. However, Codex provided a run_local_example.sh script, which improved practical usability despite the less polished written documentation. This distinction matters: a pipeline can be well described and still fail during execution, while a less polished implementation may be easier to test if it includes concrete run commands.

The recurring pattern was that agents were better at producing the shape of a pipeline than at preserving the exact semantics of each stage. For example, several outputs contained sensible directory names, modules, and MultiQC calls, yet later failed because a process expected a file name that was not actually emitted, a channel contained a stream object where a path was expected, or a parameter was not correctly passed into a process. These are not exotic failure modes; they are common when a Nextflow pipeline is assembled without end-to-end tests. The important point is that they are difficult for a non-specialist to diagnose from the final error message alone.

### Setup and execution failures revealed different debugging burdens

Each generated pipeline required modifications before or during execution. Some fixes were fair consequences of the original prompt not specifying every environmental detail, such as the need to search recursively through subdirectories for FASTQ files. Other failures were more fundamental. Biomni generated a final MultiQC process that failed with a DataflowStreamReadAdapter path-value error, missed expected raw MultiQC outputs, and had problems locating the Bowtie2 index. Claude Code failed immediately when the help command encountered an invalid label invocation, later produced a null process-input declaration for min_length, and emitted a cutadapt error after passing null where an integer was expected. Codex lacked a help option, required similar FASTQ-discovery and Picard invocation fixes, and encountered a chromosome-size mismatch indicating disagreement between BAM header contig names and the chrom_sizes file. Thread propagation was a shared practical defect. All three agent-generated workflows were expected to run with eight cores, but Bowtie2 alignments ran with one core unless corrected. This kind of error can be easy to miss because the pipeline still runs and produces files; the failure appears as unexpectedly long runtime rather than an obvious crash. For production use on an HPC system, resource propagation should be treated as a testable requirement, not an implementation detail.

### Final reports did not reproduce the merged-group reporting requirement

The most important biological reporting failure was shared by all three generated pipelines. The hand-coded pipeline produced a MultiQC report containing peak annotations, profile plots, and heatmaps for the merged groups. The agent-generated pipelines performed merging by group as requested but failed to produce the corresponding merged-group reports. This is a critical distinction. In CUT&Tag analysis, merged group-level outputs are not merely optional summaries; they often correspond to the comparisons that researchers intend to interpret biologically. A pipeline that correctly processes individual samples but fails to report merged antibody/condition groups can appear complete while omitting the primary analysis endpoint.

This failure suggests a specific limitation of natural-language pipeline generation. The prompt explicitly requested merged-group peak calling and final reports summarizing merged sample groups. Nevertheless, the generated implementations did not consistently carry merged outputs forward into annotation, plotting, and MultiQC. The agents therefore captured a local subtask - merging and/or group-level peak calling - without preserving the full downstream contract. In workflow engineering terms, the problem was not simply that a step was missing. The problem was that a data product changed category from intermediate result to final report input, and that semantic status was not enforced by tests.

### Sample-level MultiQC reports showed incomplete agreement with the reference

For non-merged sample-level reports, none of the three generated reports fully agreed with the hand-coded reference report (**Supplemental Figure 1**). Codex was closest to the reference for primary mapped-read counts. Its mapped counts were within roughly 0.0 to 0.5 million reads of the reference primary stage, but its total-read counts were approximately 1.7 to 6.4 million reads higher. As a result, Codex reported mapped percentages of 94.6% to 97.6% rather than the reference’s 100% for already filtered reference-stage rows. Claude Code showed a mixed pattern: total reads were closest to the reference primary stage, but mapped reads were closest to the noBlacklist stage and were 7.8 to 24.2 million lower than the closest reference stage. Biomni was least aligned for mapped/read-count agreement, with mapped/read totals 10.2 to 26.4 million reads below the closest reference stage.

The read-count comparison highlights an important interpretive issue. A MultiQC report is not just a collection of numbers; it is a set of labels tying metrics to specific workflow stages. If a generated report compares mapped reads from one filtering stage to total reads from another, the resulting percentages can be internally consistent but biologically misleading. This explains why Claude Code could be broad and visually comprehensive while still being difficult to trust. The report had many sections, but the mapping summary did not clearly preserve the same stage semantics as the reference.

### Duplication rates were closer than mapping counts

Duplication metrics showed better agreement than mapping/read-count summaries. The reference report contained Picard duplication values on the noBlacklist rows. Biomni differed from the reference by approximately −2.2 to +0.1 percentage points. Claude Code differed by approximately −2.2 to +0.3 percentage points. Codex differed by approximately +0.8 to +1.0 percentage points. Thus, all three were relatively close for duplication alone, with Biomni and Claude Code slightly low for several samples and Codex consistently slightly high.

This result is useful because it shows that not all QC categories failed equally. Metrics generated directly by a standard tool and collected at a well-defined step may be easier for generated pipelines to reproduce. In contrast, metrics that require consistent stage labeling, custom aggregation, or integration across multiple process outputs are more fragile. This distinction should guide future prompt design and acceptance testing.

### Codex was the strongest overall generated report, but only partially

Considering both numerical agreement and report organization, Codex produced the strongest overall generated sample-level report. It was the closest to the reference for primary mapped-read counts and contained a strong CUT&Tag-specific organization, including consolidated sections for read retention, peak counts, FRiP, TSS enrichment, and peak annotation. This does not mean Codex reproduced the reference. It did not match the reference table structure or percentages, and it failed the merged-group reporting requirement. Rather, Codex was the least problematic of the three for the subset of metrics that could be compared directly.

Claude Code produced the most comprehensive report by raw section count. It included FastQC, cutadapt, Bowtie2 or HISAT2-related alignment information, samtools, Picard, FRiP, read retention, and numerous peak annotation sections. However, its mapped-read numbers disagreed substantially with the reference, so section breadth did not translate into trustworthiness. Biomni produced a more comprehensive report than the reference in terms of including FastQC, cutadapt, samtools, Picard, FRiP, mitochondrial fraction, peak counts, and read retention, but its core mapped-read agreement was poor. The reference report was narrower but correct for the audit target, consisting mainly of Picard and samtools-derived metrics at known stages.

The practical lesson is that comprehensiveness and correctness should be evaluated separately. A report with more sections may better communicate the intended scope of a workflow, but more sections also create more opportunities for stage mismatches, mislabeled metrics, and missing downstream data products. Conversely, a narrow reference report can be a better audit target when its stage definitions are unambiguous. Future evaluations should include both: a strict audit table for numerical correctness and a separate assessment of biological interpretability.

## Discussion and conclusions

This evaluation was motivated by a practical question: can a detailed natural-language prompt produce a complex, usable bioinformatics pipeline through current agentic coding systems? The answer is qualified. Biomni, Claude Code, and Codex each generated substantial outputs that would have taken time to scaffold manually. They produced plausible workflow structures, documentation, and many expected analysis steps. However, none delivered the requested workflow without significant debugging, and none produced final reports that fully agreed with the hand-coded reference. The shared failure to produce merged-group reporting is especially important because it affected the biological endpoint of the analysis, not merely software polish.

The results were disappointing in the specific sense that the pipelines generated were not ready for use by a non-programmer. A graduate student in biology could likely edit a CSV file, install software with instructions, and run a documented command.

That is not the same as diagnosing Nextflow channel-type errors, process output mismatches, null parameter propagation, stage-labeled metric discrepancies, or missing report inputs after pipeline resume. These failures require knowledge of workflow engines, sequencing analysis, file formats, and HPC execution. The generated pipelines may reduce the amount of initial typing, but they do not remove the need for a computationally skilled reviewer.

### Why plausible pipelines failed

The main failure mode was semantic drift between the prompt, the code, and the report. The agents often implemented something that resembled the requested step but did not maintain the full contract of that step through the rest of the workflow. For example, a pipeline might merge BAMs by group, but fail to feed the merged peaks into annotation, profile plots, heatmaps, and MultiQC. A pipeline might compute read counts, but label them in a way that mixes primary, noBlacklist, deduplicated, or final BAM stages. A pipeline might expose a threads parameter, but not pass it to Bowtie2. These are not failures of syntax alone. They are failures to preserve scientific intent across software boundaries.

This kind of error is particularly dangerous in bioinformatics because workflows can produce valid-looking files even when the interpretation is wrong. A BAM file can be sorted and indexed even if it represents the wrong filtering stage. A MultiQC report can be rendered cleanly even if it compares incompatible metrics. A peak file can exist even if the matched control was not the intended control group. For this reason, scientific workflow correctness should be defined in terms of data contracts and acceptance tests, not just whether processes complete.

A second failure mode was over-implementation without staged verification. The agents attempted to satisfy many interdependent requirements at once: FASTQ discovery, metadata validation, trimming, alignment, filtering, deduplication, peak calling, merging, annotation, FRiP, deepTools visualization, and MultiQC reporting. Once a workflow of this size contains several interacting errors, debugging becomes difficult because the source of a downstream failure may lie in an earlier channel transformation, a misnamed output file, an untested metadata assumption, or a report-generation step that silently receives the wrong input. A staged implementation strategy would likely have prevented some of these errors by requiring each data product to be tested before it was used by later stages.

### What the original prompt did well

The original prompt was unusually detailed compared with a typical natural-language request. It specified the expected tools, input parameters, association table columns, validation rules, output directory structure, reporting requirements, and reproducibility goals (Supplemental file 1). It also gave server-specific constraints such as working under /workdir/$USER, avoiding Docker directly on the server, using docker1 for Docker operations, preferring Apptainer or Singularity where possible, loading Nextflow 25.4.3, and not running the pipeline automatically. These details likely helped the agents produce more realistic outputs than a short prompt would have produced.

The prompt also correctly emphasized grouped CUT&Tag analysis. It required samples to be organized by species, genome, antibody, condition, replicate, control group, and merge group. It required treatment groups and control groups to be merged separately before MACS3 peak calling, and it required final reports that included merged group summaries. This made the task more representative of a real laboratory pipeline than a simple per-sample FASTQ-to-BAM workflow.

### What additional detail is needed for future prompts

The most important conclusion from this study is that detailed biological instructions are not enough. For complex pipelines, the prompt should also define executable acceptance criteria. The following additions would likely reduce debugging burden in future agent-built workflows:

#### 1. A miniature test dataset with expected outputs

The prompt should provide two to four tiny paired-end FASTQ pairs, a tiny reference genome, a matching Bowtie2 index, a chrom_sizes file, a small GTF, a TSS BED, a blacklist BED, and an association CSV. Expected row counts, stage names, and output filenames should be included. The agent should be required to run the test and pass it before declaring the pipeline complete.

#### 2. Explicit stage semantics

Each BAM stage should have a fixed name and definition: raw-aligned, primary-aligned, mitochondrial-removed, blacklist-filtered, duplicate-marked, duplicate-removed, and final. MultiQC tables should be required to report counts from specified stages only. This would directly address the mapping-percentage mismatches observed here.

#### 3. A formal sample metadata schema

The association CSV should be validated by a schema with allowed values, required columns, uniqueness constraints, boolean parsing rules, and group compatibility checks. The prompt should include examples of valid and invalid rows.

#### 4. A required data-product contract

For every final output category, the prompt should state the upstream input that must be used. For example: merged group heatmaps must use merged group BigWigs and merged group peak sets; sample FRiP must be computed against sample peaks and matched merged-group peaks; consensus peaks must be created from comparable groups only.

#### 5. Mandatory MultiQC custom-content files

The prompt should define the exact TSV or YAML files that must be emitted for read retention, peak counts, FRiP, TSS enrichment, annotation fractions, merged group summaries, and links to plots. The agent should not merely be asked to include custom sections; it should be told the file names, column names, and example rows.

#### 6. Resource propagation tests

The prompt should require a test or log check showing that the input --threads reaches Bowtie2, samtools, cutadapt, deepTools, and MultiQC where relevant. This would have caught the shared one-core alignment issue.

#### 7. Container and HPC constraints as tests

Site-specific commands such as the Picard invocation should be specified as parameters, and a dry-run or config-summary command should print the resolved executable paths. If Docker is not allowed, the generated documentation should not include Docker-only run commands for that environment.

#### 8. Required help, dry-run, and validation modes

Every generated pipeline should include --help, --validate_only, and a local example script. The help text should list all parameters and defaults. The validation mode should check file existence, Bowtie2 index prefix completeness, contig-name compatibility, association-table consistency, and expected output writability.

#### 9. Negative tests

The prompt should require tests that deliberately fail for missing R2 files, duplicated sample IDs, unmatched control_group_id values, incompatible merge_group_id rows, and chromosome-name mismatches. These tests are as important as a successful toy run.

#### 10. A final audit checklist

The agent should be required to produce a checklist mapping each prompt requirement to a file, process, output path, and test result. This would make omissions, such as missing merged-group reports, easier to detect immediately.

#### 11. A staged implementation plan with test gates

The agent should be required to produce a development plan before writing the full workflow. That plan should divide the task into small increments: FASTQ discovery and metadata validation; raw and trimmed QC; alignment; filtering and deduplication; signal-track generation; per-sample peak calling; merged-group peak calling; annotation and FRiP; deepTools visualizations; and final MultiQC reporting. Each increment should have a specific command, expected files, and pass/fail tests. The agent should not proceed to the next stage until the current stage has passed its tests.

#### 12. Reusable agent skills for implementation and testing

Where the agent platform supports reusable skills, the prompt should instruct the agent to use them explicitly. A feature-development skill can enforce the discipline of planning, implementing, testing, and documenting each feature before moving on. Separate testing skills could define how to build toy datasets, write Nextflow smoke tests, inspect MultiQC custom-content files, compare stage-level metrics to a reference table, and verify that merged treatment and control groups are propagated into downstream reports. Skills are useful because they turn recurring expert procedures into checklists that the agent can reapply, rather than relying on a single large prompt to encode every engineering habit.

These additions shift the task from ‘write me a pipeline’ to ‘write a pipeline that proves it satisfies this contract.’ That shift is probably necessary for agentic systems to be useful in production bioinformatics. It also changes the role of the human user. Instead of reviewing thousands of lines of generated code directly, the human reviews the specification, acceptance tests, and final audit table. This is a more realistic path for collaboration between domain experts and coding agents.

They also suggest that prompt quality should be evaluated partly by how well it constrains the development process, not only by how much biological detail it contains. A prompt that asks for a complete pipeline in one response may produce an impressive repository, but it gives the agent too many opportunities to create untested assumptions. A better prompt is closer to a software development protocol: define the contract, implement one component, test it, record the result, and then extend the workflow only after the previous component is correct.

### Recommendations for using coding agents in bioinformatics pipeline construction

Based on this evaluation, coding agents are best used as accelerators under expert supervision rather than autonomous pipeline authors. A practical workflow would be to ask the agent to generate a narrow scaffold, run a tiny test dataset, add one pipeline stage at a time, and enforce tests after each stage. The user should resist the temptation to request a full production workflow in one pass unless the prompt includes a test harness and strict output contract. Large prompts can create impressive initial code, but they also increase the number of interacting requirements that may be partially implemented and hard to debug.

In practice, the first agent task should be planning rather than coding. The agent should be asked to write an implementation plan that lists the stages, expected inputs and outputs, validation checks, test commands, and criteria for moving to the next stage. The plan should then be executed in order. For example, the first deliverable might only be recursive FASTQ discovery plus association-table validation, with tests for paired-end matching and genome filtering. Only after that passes should the agent add FastQC and cutadapt. Alignment should be added only after trimmed FASTQs are verified. Filtering should be added only after aligned BAMs and alignment statistics are correct. Merged-group peak calling should be added only after sample-level peak calling has passed. Reporting should be the final layer, not a parallel task attempted before the data products it summarizes have been validated.

This staged approach is especially important for Nextflow because many serious errors occur at the interface between processes: channel shape, tuple order, metadata propagation, process output declarations, and resume behavior. These are difficult to diagnose after the full workflow has been generated. They are much easier to detect when the pipeline is built incrementally, and each stage emits a small, inspectable set of outputs. The same principle applies to biological validation. It is easier to confirm treatment-control matching and merged-group behavior with a toy association table than to infer those relationships after a full run has already produced dozens of downstream files.

Reusable agent skills could make this staged process more reliable. A feature-development skill, for example, can prompt the agent to define the feature, identify affected files, implement the smallest useful change, run the relevant tests, update documentation, and summarize what changed. A Nextflow-testing skill could require nf-core-style linting where appropriate, a minimal local test run, inspection of .command.log and .command.err files, and explicit checks for declared outputs. A bioinformatics-QC skill could require comparison of read counts across stages, verification of chromosome naming, confirmation that control groups match treatment groups, and inspection of MultiQC custom data before the report is accepted. These skills do not guarantee correctness, but they reduce the chance that the agent will skip routine checks that experienced workflow developers perform automatically.

For laboratories, the most useful near-term role for these agents may be documentation, configuration templates, module boilerplate, helper scripts, and exploratory implementations. They can also help explain errors once a knowledgeable user identifies the relevant logs. However, the final authority should remain a validated reference workflow, known-good test data, and manual review of key biological outputs. For sequencing pipelines, ‘runs without crashing’ should never be accepted as equivalent to ‘scientifically correct.’

### Other agentic systems and future comparisons

Biomni, Claude Code, and Codex are not an exhaustive set of agentic systems. They were selected because they represent relevant categories: a biomedical AI agent, a terminal/codebase-oriented coding agent, and a general software-engineering agent. Other systems may be valuable in future comparisons. ClawBio is particularly relevant as an agent-agnostic bioinformatics skill library designed to provide versioned, reproducible skills and validated code bundles for bioinformatics tasks (Corpas, M et al. 2026). AutoBA is an autonomous AI agent for automated multi-omic analyses and includes code execution and repair elements (Zhou et al., 2024). SpatialAgent focuses on spatial biology and single-cell workflows with dynamic tool execution and adaptive reasoning (Wang H. et al. 2025). BioAgents uses a multi-agent approach to help extract methods, generate executable workflows, and support human-in-the-loop bioinformatics analysis (Mehandru et al., 2025). Browser-based systems such as Pipette.bio may also be relevant for users who want an analysis service rather than a local codebase-editing agent (https://pipette.bio/).

Future work should compare these systems on multiple pipeline tasks, not only CUT&Tag. Useful benchmark tasks would include RNA-seq differential expression, ATAC-seq peak calling and differential accessibility, single-cell RNA-seq preprocessing, variant calling, and metagenomic profiling. Each task should include a reference implementation, a small test dataset, expected outputs, a debugging log, and a blinded evaluation of report correctness. The field would also benefit from standardized “workflow construction benchmarks” in which agents are scored not only on generated code, but also on installation burden, test coverage, data-contract preservation, correctness of biological outputs, and the expertise required to fix failures.

### Limitations

This study has several limitations. First, it evaluates one complex task rather than a broad benchmark suite. The observed failures may reflect the difficulty of this particular CUT&Tag workflow, the prompt design, the execution environment, or the current versions of the agents used. Second, some assessments are subjective, especially documentation quality and the expertise required for debugging. Third, the hand-coded pipeline is treated as the reference for this comparison, but it should not be interpreted as an independent biological truth set. Fourth, the current analysis focuses on final reports and observed failures rather than a line-by-line code audit of every generated workflow. Finally, the results may change rapidly as agentic coding systems improve. These limitations argue for repeated, versioned evaluations rather than one-time claims about any tool.

### Conclusion

Agentic coding systems can now produce substantial bioinformatics workflow scaffolds from detailed natural-language prompts. In this evaluation, that capability was real but insufficient. Biomni, Claude Code, and Codex each generated plausible CUT&Tag pipelines, but all required debugging, all failed the merged-group reporting requirement, and none fully agreed with the hand-coded reference report. Codex was the best overall generated report for this task because it was closest to the reference for primary mapped-read counts and had strong CUT&Tag-specific report organization. Claude Code was the most comprehensive by section count but had problematic mapping summaries. Biomni had the strongest subjective documentation but the poorest mapped/read-count agreement. The broader conclusion is that prompt-to-pipeline development is promising, but production bioinformatics requires explicit tests, precise data contracts, stage-aware reporting, and expert oversight.

## Methods

### Pipeline construction task

Each agent was given the same detailed prompt asking for a production-quality Nextflow DSL2 pipeline for paired-end CUT&Tag data (**Supplemental File 1**). The requested workflow began with recursive FASTQ discovery and pairing validation, followed by association-table validation, raw FastQC, MultiQC aggregation, paired-end adapter trimming with cutadapt, post-trim FastQC, Bowtie2 alignment, samtools sorting and indexing, alignment statistics, mitochondrial read removal, optional blacklist filtering, duplicate removal with Picard, optional UMI-aware deduplication, read-retention summaries, normalized BigWig and BedGraph generation, per-sample MACS3 peak calling, merged group and matched control peak calling, peak annotation, FRiP calculation, deepTools profile and heatmap generation, TSS enrichment summaries, and multiple MultiQC reports.

The prompt also specified the expected association CSV columns: sample_id, species, genome, antibody, condition, replicate, group_id, is_control, control_group_id, merge_group_id, peak_calling_mode, and notes (**Supplemental Table 1**). Required validation included matching FASTQ-derived sample IDs to association rows, restricting processing to samples whose genome matched the --genome parameter, verifying group compatibility, and failing early for missing or ambiguous treatment-control relationships unless control-free peak calling was explicitly allowed.

**Table 1.**
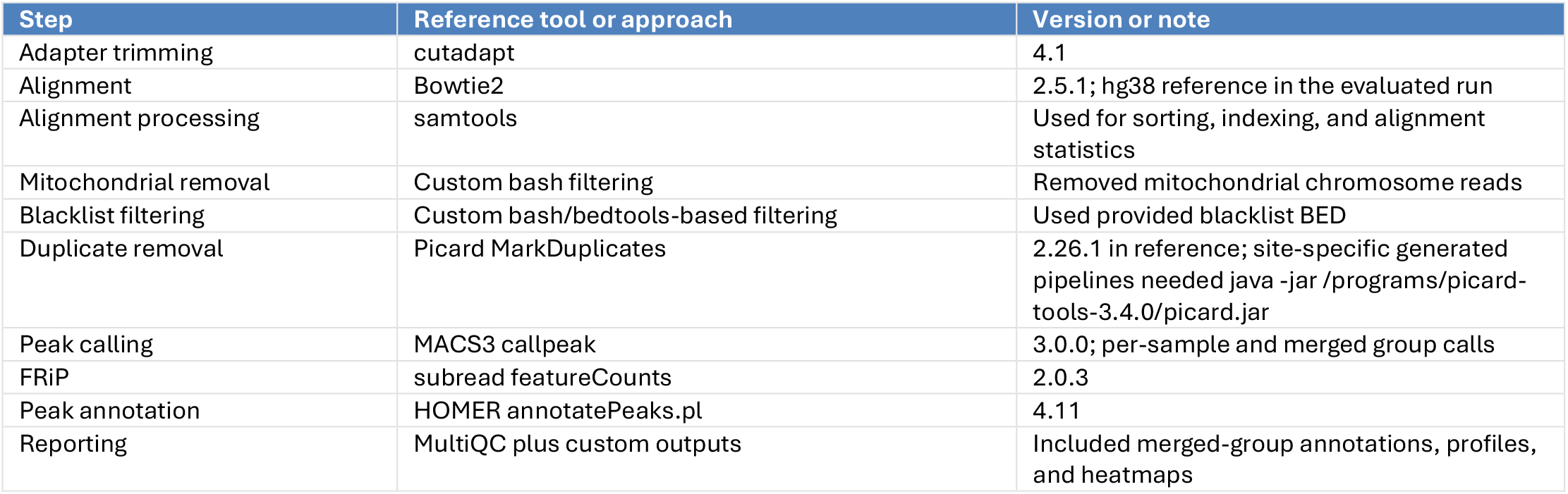
Reference CUT&Tag pipeline steps, tools, and software versions. Summary of the tools and approaches used in the hand-coded reference CUT&Tag pipeline against which the agent-generated pipelines were evaluated. The workflow begins with adapter trimming using cutadapt, followed by paired-end alignment with Bowtie2 to the hg38 reference genome in the evaluated run. Alignment files were sorted, indexed, and summarized with samtools, then filtered to remove mitochondrial reads and reads overlapping blacklisted genomic regions. Duplicate fragments were removed with Picard MarkDuplicates, and peaks were called with MACS3 for both individual samples and merged treatment groups. FRiP scores were calculated using subread featureCounts, and peak annotations were generated with HOMER annotatePeaks.pl. Final reporting used MultiQC together with custom outputs, including merged-group annotations, enrichment profiles, and heatmaps. The table also records relevant version information and implementation notes, including the site-specific Picard invocation required for some generated pipelines.

| Step | Reference tool or approach | Version or note |
| --- | --- | --- |
| Adapter trimming | cutadapt | 4.1 |
| Alignment | Bowtie2 | 2.5.1; hg38 reference in the evaluated run |
| Alignment processing | samtools | Used for sorting, indexing, and alignment statistics |
| Mitochondrial removal | Custom bash filtering | Removed mitochondrial chromosome reads |
| Blacklist filtering | Custom bash/bedtools-based filtering | Used provided blacklist BED |
| Duplicate removal | Picard MarkDuplicates | 2.26.1 in reference; site-specific generated pipelines needed java -jar /programs/picard-tools-3.4.0/picard.jar |
| Peak calling | MACS3 callpeak | 3.0.0; per-sample and merged group calls |
| FRiP | subread featureCounts | 2.0.3 |
| Peak annotation | HOMER annotatePeaks.pl | 4.11 |
| Reporting | MultiQC plus custom outputs | Included merged-group annotations, profiles, and heatmaps |

### CRAFT prompt construction

The task prompt was written using the CRAFT framework: context, role, action, format, and target audience. The context described the agent as a biomedical AI system expected to combine LLM reasoning, retrieval-augmented planning, tool selection, and executable code generation. The role framed the model as a senior computational biologist and workflow engineer. The action section listed the required pipeline behavior in detail. The format section required a complete blueprint and implementation package, including main.nf, nextflow.config, modules, helper scripts, environment specifications, and README content. The target audience was a computational biology researcher or bioinformatics engineer who could edit CSV files and run command-line tools but needed clear documentation.

The CRAFT structure was intended to reduce ambiguity and make the task closer to a realistic project specification. It was not sufficient by itself

### Generated pipeline execution and debugging assessment

Each generated pipeline was set up, modified as needed, and run against the same FASTQ inputs. Problems encountered during setup and execution were recorded qualitatively. Bugs were grouped by likely cause, including missing functionality, to guarantee correctness. Based on the results, future CRAFT prompts for workflow generation should include executable acceptance tests, exact output contracts, and expected metric values from a minimal test dataset (**Supplemental Figure 2**).

### Hand-coded reference pipeline

The reference pipeline was written by the authors and used as the comparison baseline. It processed CUT&Tag FASTQ files through adapter trimming with cutadapt version 4.1, alignment to the reference genome with Bowtie2 version 2.5.1, mitochondrial read removal, blacklist filtering using a provided BED file, duplicate removal with Picard version 2.26.1, per-sample peak calling with MACS3 version 3.0.0, group-level peak calling with MACS3 version 3.0.0 after merging treatment and matched control groups, FRiP calculation with featureCounts version 2.0.3, and peak annotation with HOMER version 4.11. The workflow also generated signal tracks, QC summaries, and MultiQC reports (**Table 1**).

The reference pipeline included custom logic to group samples by species, genome, antibody, condition, and merge_group_id. Samples sharing a treatment merge group were merged into group-level BAM files. Control BAMs were merged according to the corresponding control_group_id. MACS3 was then run on each merged treatment group against its matched merged control group. The reference also produced group-level peak sets and downstream reporting for merged groups, including peak annotations, profile plots, and heatmaps.

environment mismatch, parameter propagation, Nextflow syntax or channel errors, missing expected outputs, metadata association problems, and reporting failures. The assessment also recorded whether an error would likely be fixable by a wet-lab researcher following documentation or whether it required deeper programming or Nextflow knowledge.

All three pipelines required a modification to search for FASTQs in subdirectories. This was considered a fair limitation because the initial prompt did not make that behavior explicit enough, although the final specification did call for recursive input scanning. All three also required correction of the Picard invocation in the tested server environment. This was treated as an environment-specific issue but also as evidence that tool invocation should be parameterized and validated.

### MultiQC report comparison

Final MultiQC reports from the generated pipelines were compared with the hand-coded reference report. The comparison focused on the presence or absence of requested report sections, agreement of sample-level mapping and read-count metrics, duplication-rate agreement, and the presence of merged-group reporting. For mapping/read-count comparisons, generated sample-level flagstat rows were compared against the closest matching reference stage. Because not all reports used identical stage labels, the comparison explicitly recorded which reference stage was closest for each generated report. Duplication values were compared against Picard duplication values on the reference noBlacklist rows.

Comprehensiveness was assessed separately from numerical agreement. A report could therefore be classified as more comprehensive than the reference while still disagreeing with it numerically. This separation was necessary because the reference report was intentionally treated as the audit target rather than the maximal possible biological QC report.

## Data and code availability

The code repositories for the evaluated agent-generated pipelines are listed below.

- Biomni-generated pipeline: https://github.com/paulmunn/Biomni_CUTnTag_pipeline
- Claude Code-generated pipeline: https://github.com/paulmunn/Claude_CUTnTag_pipeline
- Codex-generated pipeline: https://github.com/paulmunn/Codex_CUTnTag_pipeline
- Supplementary File 1: full CUT&Tag pipeline construction prompt used for all three agents.

## Acknowledgments

We thank Faraz Ahmed of Cornell University’s Transcriptional Regulation & Expression Facility for his assistance in the development and refinement of the hand-coded CUT&Tag pipeline used as the reference workflow for this study. We also thank the Cornell University Genomics Facility for providing access to unpublished sequencing data used to test and evaluate each of the pipelines examined in this study.

## Supplemental Figures and Table

**Supplemental Figure 1.**
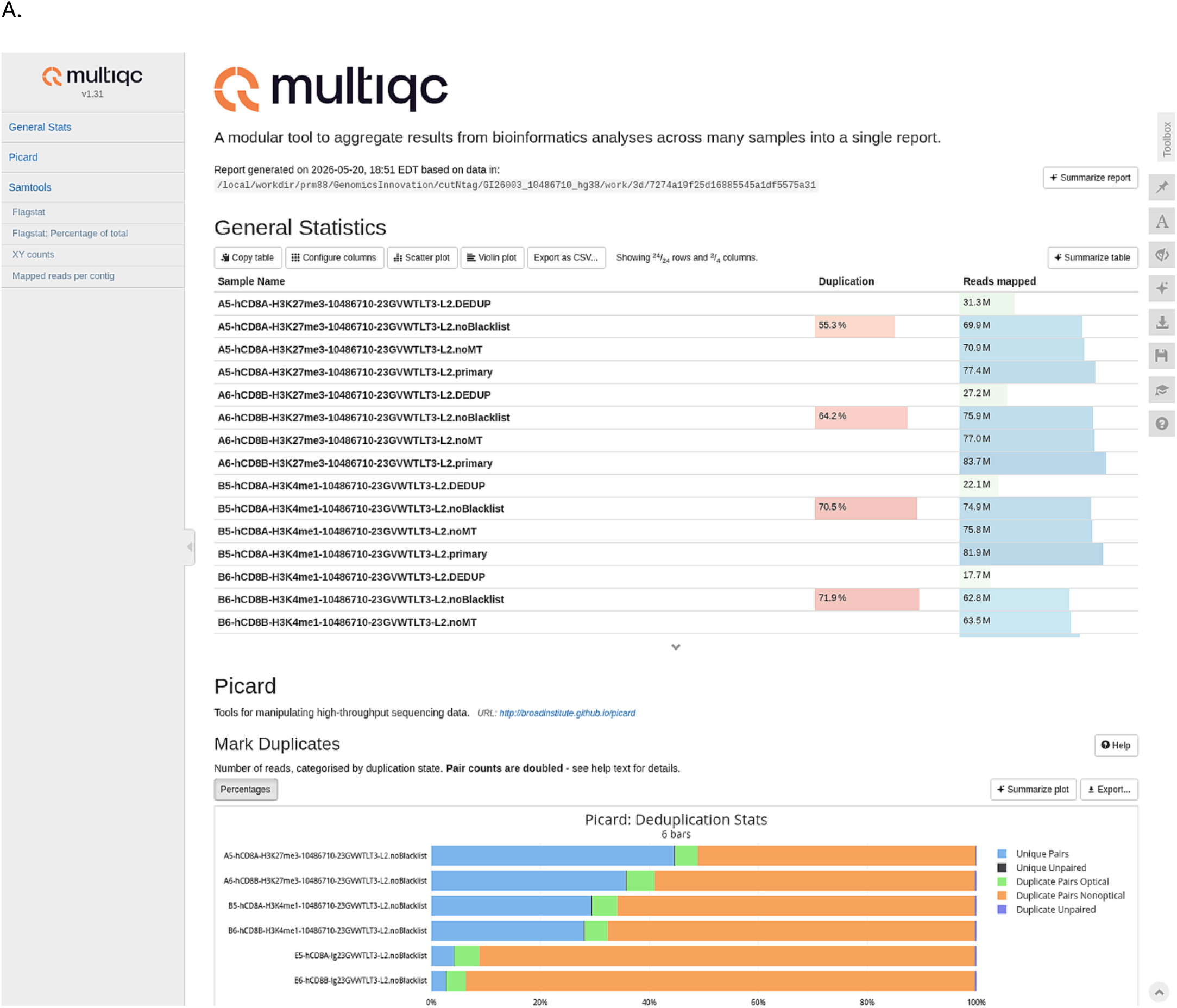
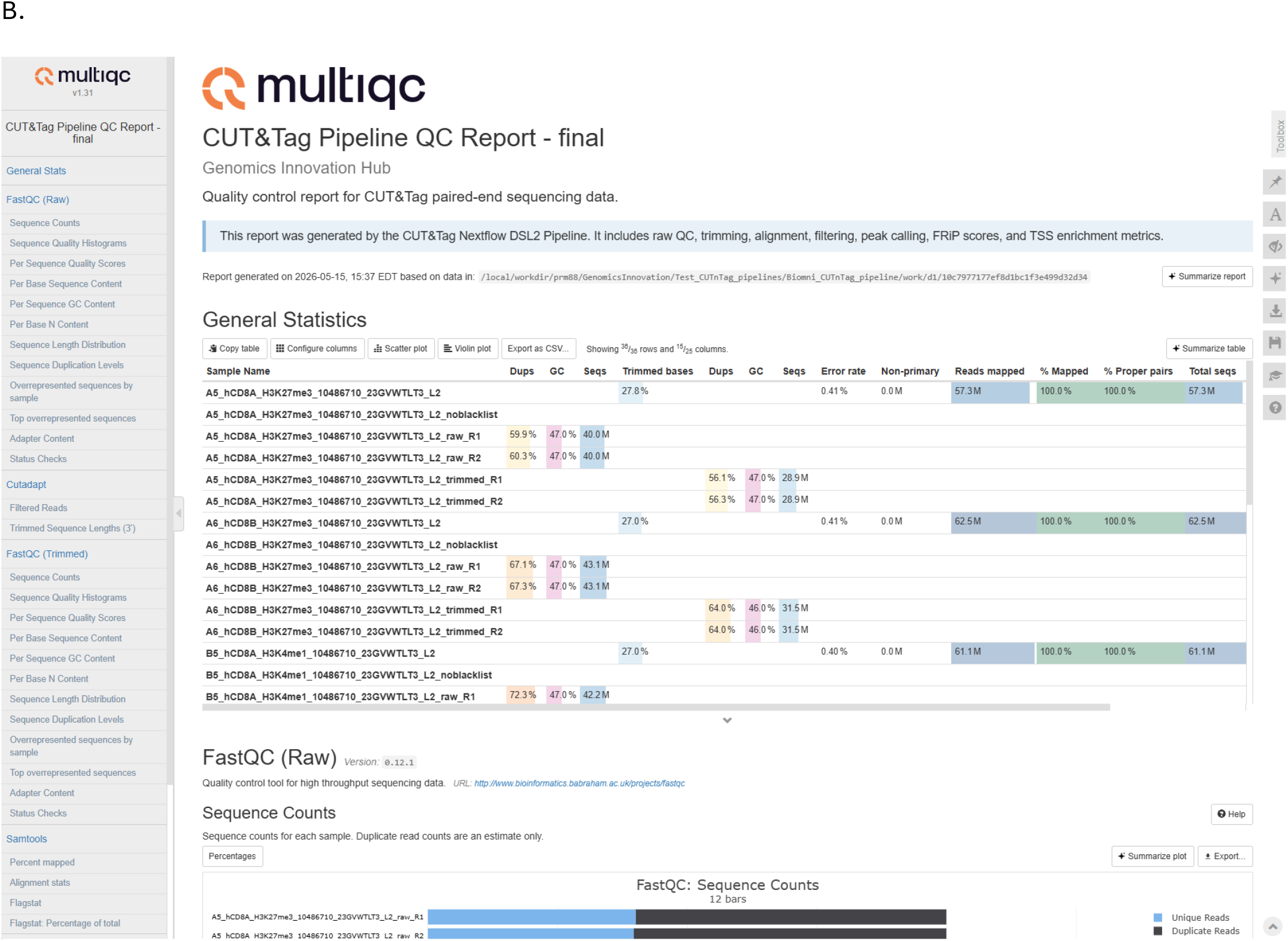
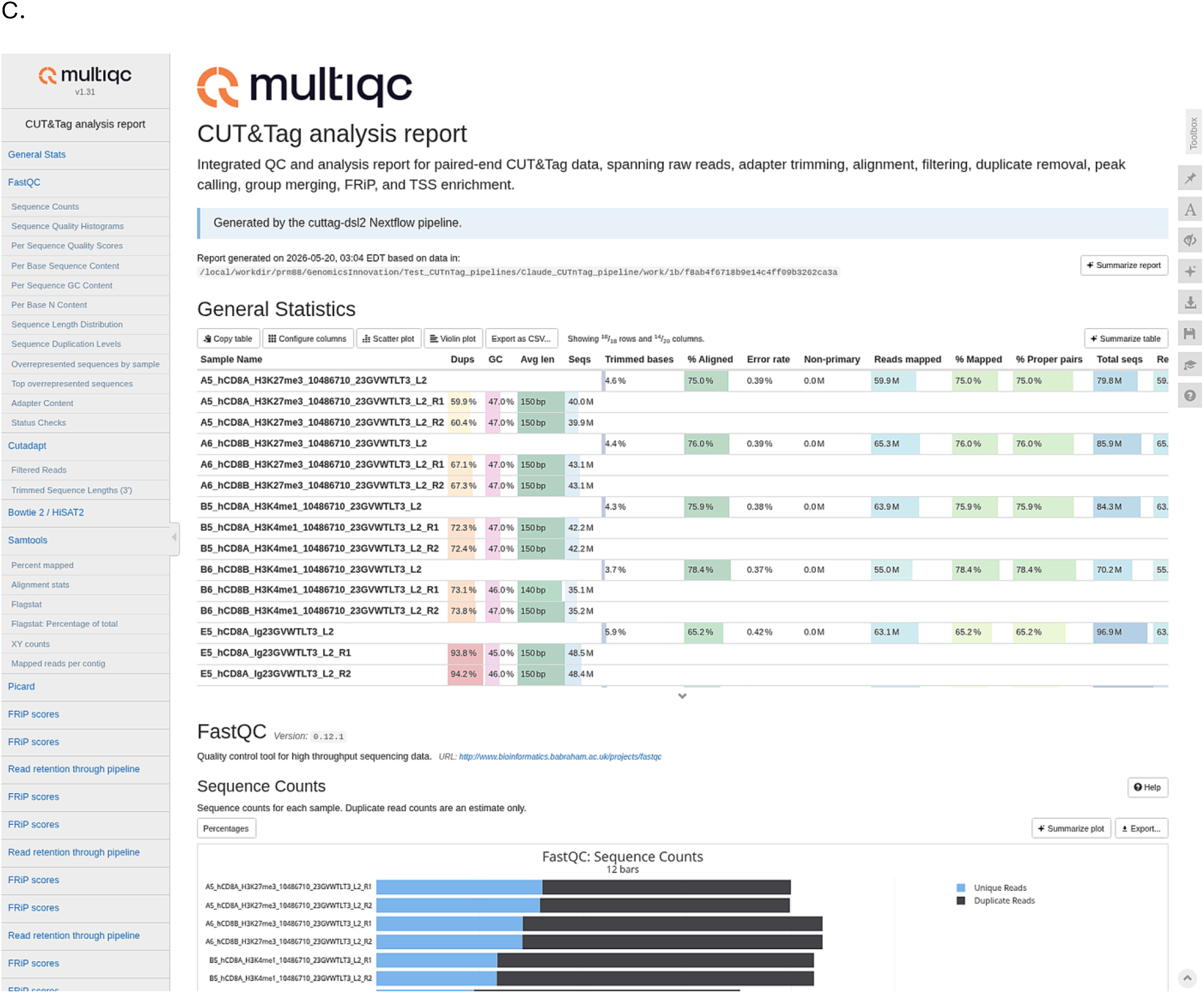
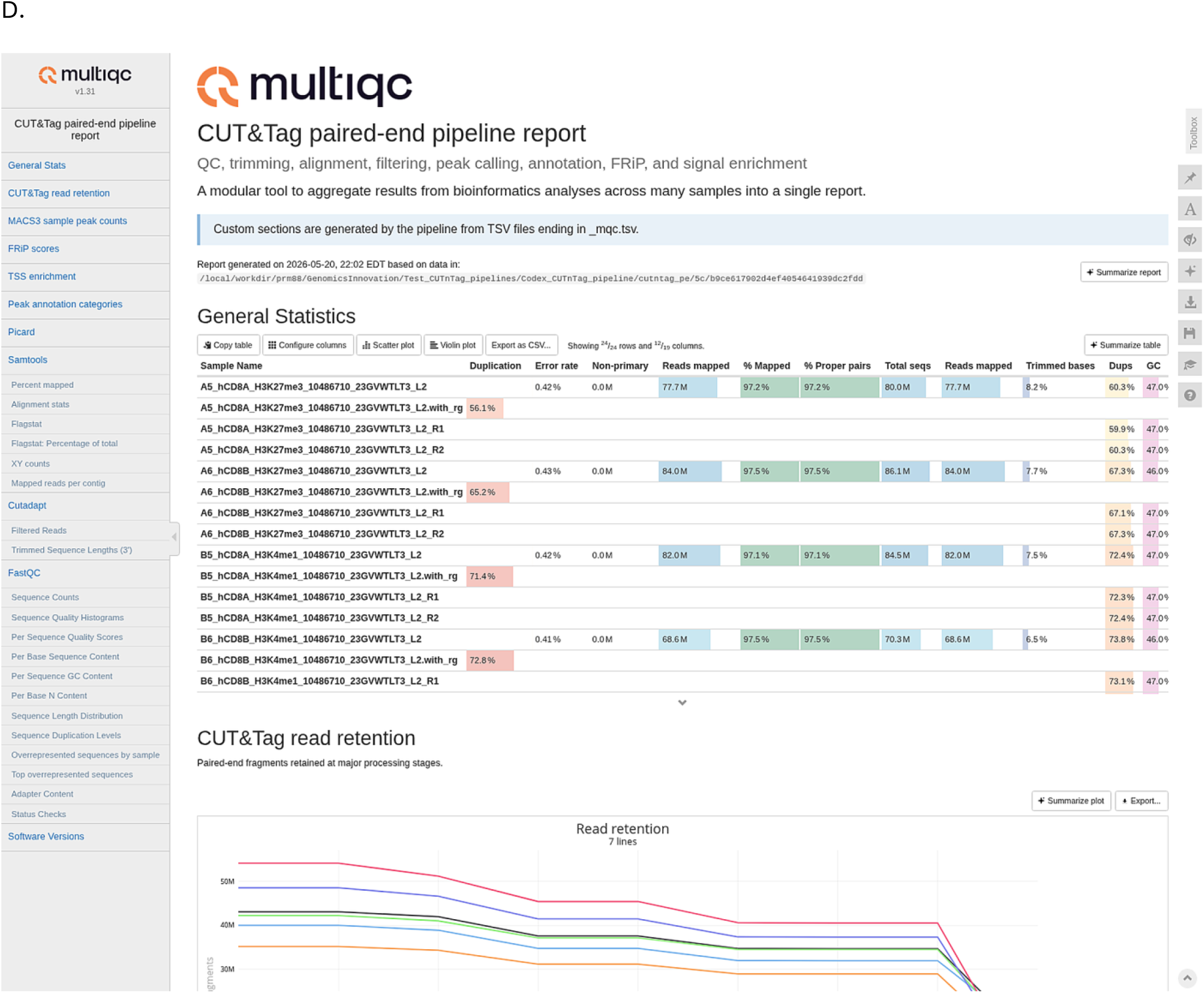
Representative screenshots from final QC reports generated by the reference and AI-produced CUT&Tag pipelines. Representative panels from the final MultiQC reports are shown for the (A) **hand-coded reference pipeline**, (B) **Biomni**, (C) **Claude Code**, and (D) **Codex**. The screenshots were selected to emphasize comparable report regions, allowing visual comparison of how each pipeline summarizes core CUT&Tag quality-control outputs and report organization. The figure illustrates differences in report structure, section completeness, and the presentation of key summary metrics across implementations. In the reference pipeline, the report serves as the ground-truth benchmark for downstream comparison. The AI-generated reports vary in both scope and layout, with differences in the inclusion and organization of biologically relevant CUT&Tag summary sections such as mapping and filtering metrics, read-retention summaries, peak-related outputs, FRiP, TSS enrichment, and annotation summaries.

**Supplemental Figure 2.**
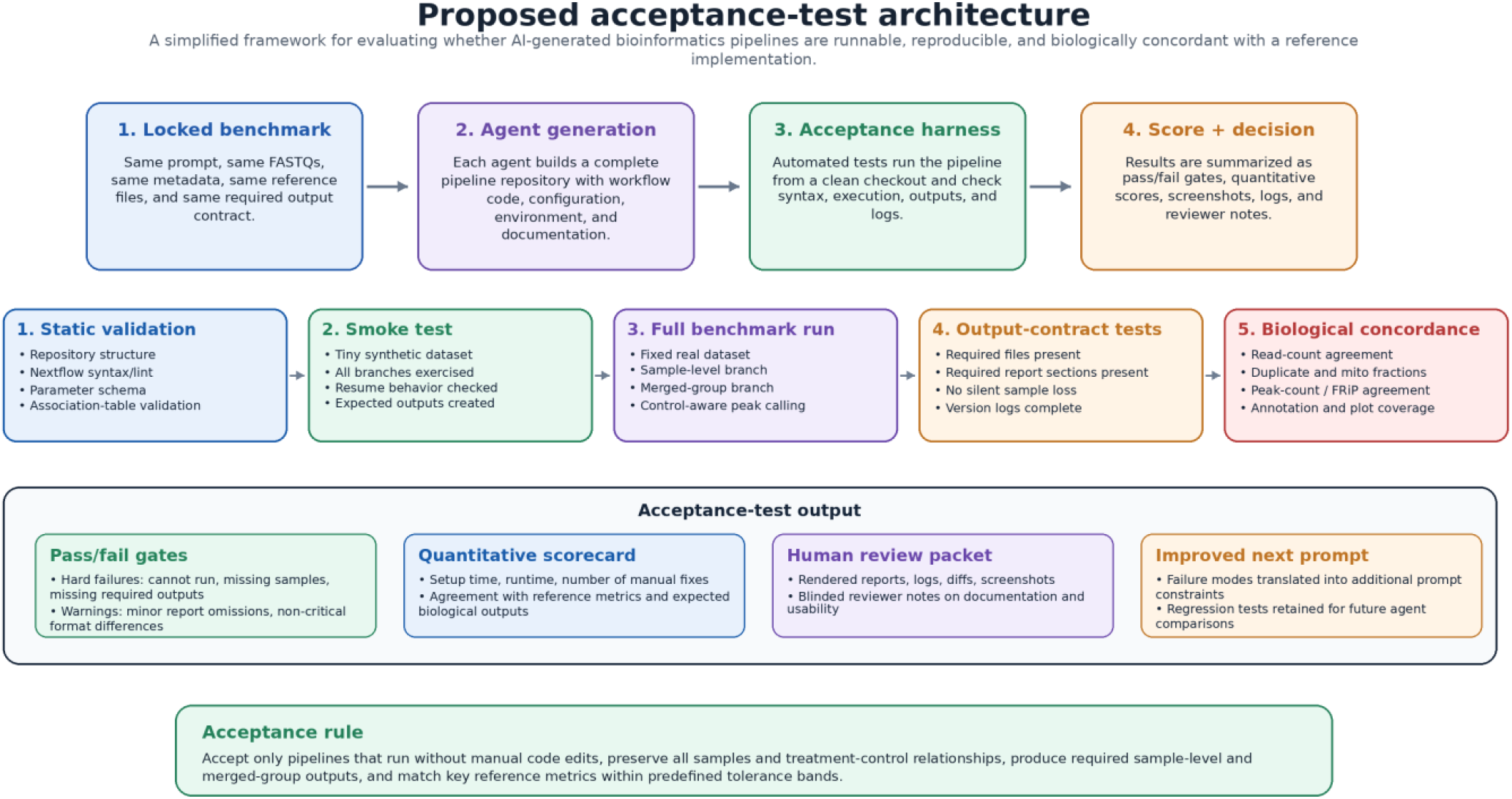
Proposed acceptance-test architecture for future prompt-to-pipeline evaluations. The proposed framework separates agent-based pipeline generation from formal acceptance testing. Each coding agent receives the same locked benchmark package, including the task specification, test datasets, reference files, expected outputs, execution contract, scoring rubric, and audit metadata. The resulting repositories are then evaluated through a standardized test harness rather than by ad hoc manual inspection. The acceptance process begins with static validation of repository structure, Nextflow syntax, parameter schemas, and metadata tables, followed by a smoke test on a minimal dataset to confirm that all major workflow branches execute. Pipelines that pass this stage are run on a fixed benchmark dataset and assessed for required output files, report sections, sample retention, treatment-control matching, merged-group behavior, and reproducibility metadata. Biological concordance is then evaluated by comparing read-count summaries, filtering metrics, duplicate and mitochondrial fractions, peak counts, FRiP scores, annotation summaries, and merged-group outputs against a hand-coded reference implementation. Final outputs from the framework include hard pass/fail gates, a quantitative scorecard, a human-review packet, and a record of failure modes that can be incorporated into future prompts and regression tests. The goal is to make future prompt-to-pipeline comparisons reproducible, auditable, and less dependent on subjective debugging impressions.

**Supplemental table 1.** Association table design for treatment-control mapping in the CUT&Tag workflow. Example association table used to define sample metadata, biological grouping, and treatment-control relationships for the CUT&Tag pipeline. Each row represents one FASTQ-derived sample and records the species, genome build, antibody target, replicate number, biological group identifier, and whether the sample is a control. Treatment samples targeting H3K27me3 or H3K4me1 are assigned to antibody-specific groups, such as hCD8_H3K27me3 or hCD8_H3K4me1, while the IgG samples are assigned to the control group hCD8_Ig23GVWTLT3. The Control Group ID column links each non-control treatment sample to the appropriate matched IgG control group used for control-aware peak calling. Control samples are marked as TRUE in the Is_Control column and do not require a control-group assignment. This metadata structure allows the pipeline to validate sample identities, process only samples matching the selected genome, merge biological replicates by group, merge matched controls, and run MACS3 peak calling for each treatment group against the correct control group.

| Sample ID | Species | Genome | Antibody | Replicate | Group ID | Is_Control | Control Group ID |
| --- | --- | --- | --- | --- | --- | --- | --- |
| A5_hCD8A_H3K27me3_10486710_23<br>GVWTLT3_L2 | human | hg38 | H3K27me3 | 1 | hCD8_H3K27me3 | FALSE | hCD8_Ig23GVWTLT3 |
| A6_hCD8B_H3K27me3_10486710_23<br>GVWTLT3_L2 | human | hg38 | H3K27me3 | 2 | hCD8_H3K27me3 | FALSE | hCD8_Ig23GVWTLT3 |
| B5_hCD8A_H3K4me1_10486710_23G<br>VWTLT3_L2 | human | hg38 | H3K4me1 | 1 | hCD8_H3K4me1 | FALSE | hCD8_Ig23GVWTLT3 |
| B6_hCD8B_H3K4me1_10486710_23G<br>VWTLT3_L2 | human | hg38 | H3K4me1 | 2 | hCD8_H3K4me1 | FALSE | hCD8_Ig23GVWTLT3 |
| E5_hCD8A_Ig23GVWTLT3_L2 | human | hg38 | IgG | 1 | hCD8_Ig23GVWTLT3 | TRUE |  |
| E6_hCD8B_Ig23GVWTLT3_L2 | human | hg38 | IgG | 2 | hCD8_Ig23GVWTLT3 | TRUE |  |

